# Optogenetic Analysis of Behavior in the Mosquito *Aedes aegypti*

**DOI:** 10.64898/2026.03.15.711871

**Authors:** Spruha Rami, Michelle So, Cassandra Travis, Yaoyu Jiao, Paul Shamble, Trevor R. Sorrells

**Author notes:** Equal contribution, listed alphabetically (Spruha Rami, Michelle So, Cassandra Travis). Correspondence should be addressed to T.R.S.

## Abstract

The mosquito Aedes aegypti is an important vector of viral pathogens and serves as a model for other vector species. Pathogens are transmitted when a mosquito bites a host animal, but the neural circuits that control seeking and biting behavior are not known. Here, we detail methods for the manipulation of neural activity in the mosquito using optogenetics, a key technique to determine the causal relationship between neural circuits and behavior. These methods include rearing mosquitoes for optogenetics and three assays that are designed to measure different steps in the sequence of arousal, attraction, proboscis probing, and engorgement on the blood of host animals. These behaviors occur at different spatial scales and in response to different sensory stimuli. Each behavioral assay is outfitted with red (~625 nm) LEDs for optogenetic activation. To detect arousal in response to olfactory stimuli, flight and walking are quantified. To assay attraction or landing, mosquitoes are presented with a heated blood meal in a large arena. Proboscis probing and engorgement are assayed with video resolution that enables measurement of appendages and abdomen size. The protocol describes machine vision models to enable high-resolution temporal quantification of behavior as well as endpoint measurements of feeding. These methods can be used to test the role of any genetically accessible population of neurons in mosquito biting behavior and can be extended to additional behaviors.

## Introduction

The mosquito is one of the most dangerous animals in the world, not for its ferocity or strength but for its efficiency as a vector for disease. Malaria-causing parasites spread by *Anopheline* mosquitoes are responsible for hundreds of thousands of deaths yearly, while *Aedes aegypti* and *Aedes albopictus* are responsible for the spread of yellow fever, dengue, chikungunya, Zika fever, and many other life-threatening diseases^1^. Mosquitoes are present on every continent except Antarctica and pose a great concern for public health^2^. Blood-feeding behavior directly transmits pathogens, but it is also an essential step in the mosquito life cycle, as a vertebrate blood meal is required to lay eggs. Curbing the spread of mosquito-borne diseases thus requires understanding their host-seeking mechanisms.

It is well-established that female mosquitoes make use of long- and short-distance cues to locate hosts^3,4^. Carbon dioxide (CO_2_) is a highly volatile gas present in human exhalation and is often the first cue that a mosquito senses. It is detected in *Aedes aegypti* olfactory sensory neurons by Gr1, Gr2, and Gr3 proteins, which form the CO_2_ receptor^5^. Exposure to CO_2_ causes take-off and flight behavior, also known as activation^6,7^, which can persist for up to 10-15 minutes, even in the absence of further stimuli^8^. CO_2_ synergizes with skin odor to enhance activation and attraction^5,9^. Following CO_2_ and skin odor detection, the mosquito is primed to sense additional, more proximal host cues, such as heat, visual contrast, and humidity that guide her to land on a viable blood meal host^5,10–13^. Integration of multiple sensory cues allows mosquitoes to effectively pursue and locate their host.

When a female mosquito lands on a potential host, she thrusts her proboscis into the surface until a patch of skin is located, a process known as probing^14,15^. Following probing, her stylet pierces the skin, locates a blood vessel, and simultaneously ingests blood while injecting saliva, doubling her body weight in several minutes^16^. Blood is directed to the midgut, where proteins are digested for egg production. Interestingly, engorgement on a host is not necessary for pathogen spread, as probing alone can transmit disease^16,17^. Therefore, understanding each of the steps in biting behavior could allow the development of strategies to disrupt female mosquito host seeking and biting and ultimately to control the spread of mosquito-borne diseases.

Numerous tools have been created in model organisms for manipulating neuronal activity. Early tools worked constitutively or by thermal activation^18–21^. While useful, these genetic reagents pose certain drawbacks. Constitutive tools can affect development, leading to phenotypes not due solely to the circuit function of the neurons. Mosquitoes use body heat to identify host vertebrates, making thermally activated channels uniquely unsuitable for studies of their behavior^22,23^.

In studies of behavior, optogenetics refers to the use of genetic tools that respond to specific wavelengths of light to control the behavior of neurons of interest. This technique has already been used in multiple systems, from examining the valence of thirst in mice, to studying courtship behavior in *Drosophila*, to restoring vision loss from retinitis pigmentosa in human trials^24–26^. Optogenetic channels work by absorbing photons and allowing specific types of ions to pass through the plasma membrane, activating or inhibiting the neuron. By introducing transgenic light-sensitive protein-coding genes into an organism, this technique can be used to activate or inhibit specific neurons that would otherwise be impossible to manipulate. Optogenetics provides advantages in temporal and spatial control, and a range of controls that enable experimental inference.

We recently described the first use of optogenetic tools to study mosquito behavior^8^. These tools rely on the red-light-activated cation channel CsChrimson, a red-shifted channelrhodopsin^27^. Microbial opsins, including CsChrimson, require a chemical cofactor, all-*trans* retinal, in order to absorb photons^28^. CsChrimson is fused to the tdTomato fluorescent reporter and is under the control of the QF2/QUAS binary expression system, enabling it to be flexibly combined with different driver lines^29^. To target CO_2_ sensory neurons, a driver line was used that expresses the QF2 transcription factor in neurons that express the Gr3 subunit of the CO_2_ receptor^30^. Mosquitoes showed minimal behavioral response after exposure to red light alone^8^; additionally, red light is most effective at penetrating biological tissue, making a red-light-activated opsin ideal for optogenetics^25^.

Here, we describe protocols for three assays that measure different aspects of host-seeking and blood-feeding behavior combined with optogenetic stimulation (**Figure 1A**). First, we provide a protocol for the rearing of optogenetic mosquito lines. The second section describes the opto-thermocycler, a modified PCR thermocycler that can be used to track arousal and probing when mosquitoes are exposed to temperature and light stimuli. The third section describes the blood blanket assay, where mosquitoes are presented with constant access to a thin artificial blood meal on the base of a modified thermocycler, enabling tracking of flight, probing, and engorgement under exposure to varying light and changing temperature conditions. Finally, we describe the opto-membrane feeder, where mosquitoes are presented with a warm blood meal while in a canister, allowing tracking of both attraction to a blood source and engorgement under exposure to light stimuli. In addition to the expected outputs for each of these assays, machine vision models are described to track the body parts of the mosquitoes and identify different behaviors performed during the assay. The opto-thermocycler, blood blanket, and opto-membrane feeder assays offer ways to measure distinct stages of mosquito blood-feeding behavior while manipulating neural activity with light. These innovative techniques enable the study of how mosquito neural circuits control behaviors that threaten public health.

**Figure 1:**
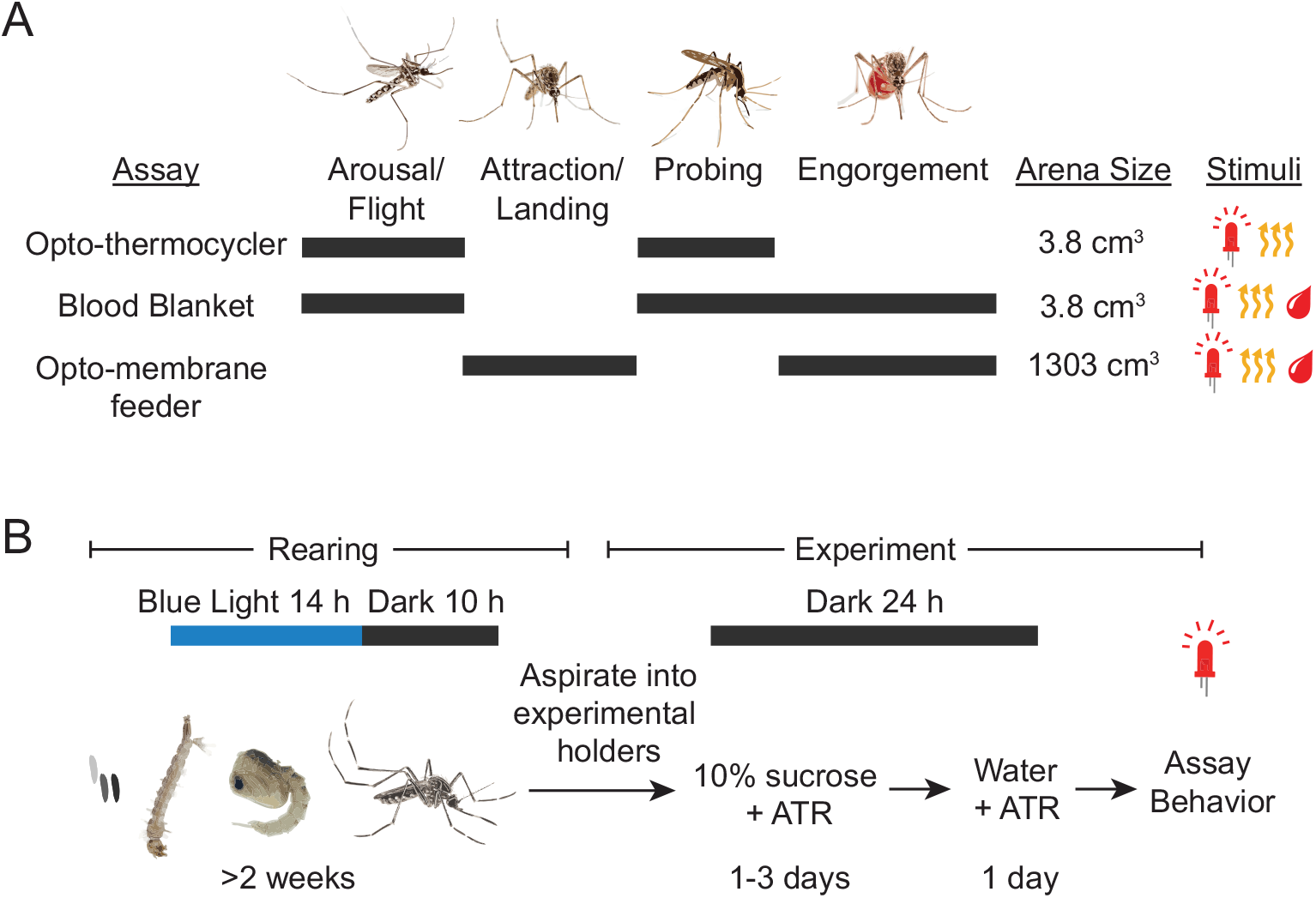
Assays and rearing for optogenetics in mosquitoes. **(A)** Schematic of mosquito host-seeking behaviors with characteristics of the behavior assays presented. The steps of host seeking that are measured by each of the behavior assays are indicated with gray bars. **(B)** Timeline of rearing of mosquitoes for optogenetics experiments.

### Protocol

1. **Optogenetic rearing**
  1.1. Rear mosquitoes according to standard protocol^31^ with the following exceptions to maximize response of optogenetic lines during experimentation.
    1.1.1. **NOTE:** Mosquitoes used in this protocol contain transgenes and under United States Department of Agriculture regulations must be reared in Arthropod Containment Level II facilities^32^. Mosquitoes are contained to prevent escape and disposed of in biohazard waste.
  1.2. Make crosses using homozygous Gr3-QF2, QUAS-CsChrimson, and wildtype Liverpool strains to generate the heterozygous experimental lines Gr3-QF2/+, QUAS-CsChrimson/+, and Gr3>CsChrimson. Mosquito genotype can be confirmed by fluorescent markers (3xP3-dsRed for Gr3-QF2 and 3xP3-CFP for QUAS-CsChrimson) or by PCR for the inserted construct^8^.
  1.3. Raise experimental animals inside of a light-tight incubator set to 28 °C and 80% humidity, on a 14 h blue light (450 nm), 10 h dark cycle (**Figure 1B**).
    1.3.1. **NOTE:** The use of blue light minimizes activation of red-light activated channels, such as CsChrimson. All rearing steps occurring outside of the incubator should be performed in low light conditions.
  1.4. Begin experiments on adult mosquitoes 1 week post eclosion at the earliest. While the ideal range for behavioral experimentation is 2-3 weeks, mosquitoes up to 4 weeks post eclosion can be used. Control and experimental genotypes should be the same age, recorded as the number of days post eclosion. To standardize the ages of the animals, the eggs for all genotypes should be hatched on the same day.
  1.5. Two days prior to experimentation, gently aspirate mated, non-blood-fed female mosquitoes out of the cages and anesthetize at 4 °C. Using a feather or aspirator, carefully transfer to the experimental containers according to the assay being used. Prepare a 0.2 M aliquot of all-***trans*** retinal (ATR) in dimethyl sulfoxide (DMSO), stored at −20 °C. Provide 10% sucrose and 400 µM ATR via soaked cotton wicks on the mesh of the container. Animals may feed for 1-3 days in the dark on this meal.
    1.5.1. **NOTE:** Feeding in the dark, as opposed to under blue light, prevents bleaching of the ATR **(Figure 1B)**.
  1.6. One day before experimentation, replace the sucrose feeders with new feeders containing distilled water and 400 µM ATR. Animals remain in the dark until experimentation **(Figure 1B)**.
    1.6.1. **NOTE:** The experiments should be run during the mosquitoes’ morning or evening activity peaks to maximize response rate.
  1.7. Proceed to section 2 opto-thermocycler, section 3 blood blanket, or section 4 opto-membrane feeder, depending on which assay is to be used.
2. **Opto-thermocycler**
  2.1. **Assembling components of the assay**
    2.1.1. Make the acrylic plates to contain the mosquitoes. Refer to the figure for diagrams of the assay (**Figure 2A**) and electronic components (**Figure 2B**), and an image of the acrylic plate (**Figure 2C**). See files in (https://github.com/sorrellslab/opto-thermocycler/releases/tag/v1.0.0 for templates.
      2.1.1.1. To create the walls, laser cut a 3 mm (1/8 inch) acrylic sheet into four 10 cm by 1 cm rectangles and six 7 cm by 1 cm rectangles. Affix the rectangles with acrylic glue into a five-by-three matrix, where each unit is a 17 mm-by-17 mm-by-10 mm square. The template for these pieces is in the file “14_animal_thick_v3.ai”.
      2.1.1.2. For the bottom of the chamber, cut a 1.5 mm (1/16 inch) thick acrylic sheet to 6.985 cm (2 and 3/4 inch) by 10.16 cm (4 inch) with the bottom right well removed. Cut a 1.5 mm (1/16 inch) thick acrylic to a 6.985 cm (2 and 3/4 inch) inch by 10.16 cm (4 inch) rectangle for the lid. The template for these pieces is “14_animal_thin_v3.ai”.
      2.1.1.3. Cut a UV-resistant black mesh sheet to the same size as the bottom acrylic piece (**Figure 2C**). Sandwich the mesh between the walls of the plate and the bottom piece, then glue them in place with acrylic glue. Remove the mesh in the bottom right well so the hole is unobstructed.
    2.1.2. Construct a frame that will hold the camera and LEDs (**Figure 2A**).
      2.1.2.1. Build a frame with four pillars at each corner of the thermocycler using 60.96 cm (2 ft), 25 mm square optical construction rails. Use two 22.86 cm (9 inch), 25 mm construction rails to connect the front-right pillar to the back-right pillar and the front-left pillar to the back-left pillar, 30 cm from the bench. Connect the right and left pillars in the front with a 30.48 cm (1 ft), 25 mm construction rail, elevated 24 cm off the bench, and in the back with a 30.48 cm (1 ft), 25 mm construction rail elevated 42 cm off the bench.
      2.1.2.2. Attach 1-foot-long optical posts to the construction rail on the left and right sides of the frame. Tie six 627 nm (red) LEDs connected in series 24 cm above the PCR block of the thermocycler to each post. Although the wavelength of light resulting in peak activity of CsChrimson is ~590nm, wavelengths near ~625nm balance the need to activate CsChrimson and avoid behavioral responses to light in genetic controls^27^. Tape a heat sink and lens on each LED to focus the light toward the block of the thermocycler.
      2.1.2.3. Fasten a 30.48 cm (1 ft) optical post to the top of the front two construction rail pillars. Use a right-angle clamp in the middle of the optical post to attach a 15.24 cm (6 inch) optical post facing inwards over the thermocycler block. Affix a camera with an infrared long-pass filter to the end so it is pointing down, 40 cm over the block.
      2.1.2.4. Using optical posts, create a frame that extends from the construction rail in a rectangle around the PCR block, elevated just above the surface. Affix a strip of infrared LEDs facing towards the PCR block for illumination of the mosquitoes at a right angle to the direction of the camera. The infrared illumination used for video imaging does not heat the arena measurably because the thermocycler surface is constantly held at 24 °C during the experiment.
      2.1.2.5. Drape a blackout curtain over the structure to prevent interference from ambient light.
    2.1.3. Cover the surface and sides of the PCR block with a single layer of blackout tape to create a flat, even surface with a dark background.
    2.1.4. Place the thermocouple sensor on the PCR block in the lower right corner and secure it with blackout tape. Confirm even distribution of light across the PCR block using a power meter. Adjust voltage or current on power supply to deliver light at an intensity of 12 µW/mm^2^.
    2.1.5. On the edge of the PCR block, place an infrared LED to synchronize the video with the stimulus delivery output (**Figure 2C**). Cover the surface with black tape to allow the light to be detected without creating a glare in the camera.
    2.1.6. Wire the thermocouple, synchronization LED, and red light LEDs according to the diagram in **Figure 2B**.
    2.1.7. Download the required software (SpinView (version 1.27.0.48), CoolTerm (version 2.0.1), Arduino IDE (version 1.8.12)).
    2.1.8. Download the Arduino code and thermocycler program from the GitHub repository (https://github.com/sorrellslab/opto-thermocycler) and transfer them to the computer and thermocycler, respectively. The Arduino code is programmed to deliver red light, body-temperature heat, and the combination of the two stimuli. There are seven stimulus programs where the stimulus order is randomized between them provided as resources. These can be modified to deliver different stimuli. **NOTE**: There are nine stimuli per experiment: three of heat only, three of light only, and three of heat and light simultaneously. Stimulus delivery operates by the Arduino detecting changes in the surface temperature of the PCR block via the thermocouple then delivering light stimuli at the specified times. Additional details can be found in comments within the stimulus programs.
  2.2. **Trial Preparation**
    2.2.1. Rear mosquitoes according to the optogenetic rearing protocol; see section 1 and **Figure 1B**.
    2.2.2. Two days before the experiment, transfer 14 female mosquitoes with an aspirator to a separate container (i.e., a cage). Provide ATR to the mosquitoes according to step 1.5. If multiple genotypes are being compared, blind them to the experimenter. **NOTE**: Once mosquitoes receive ATR, they should be handled under dim light conditions.
    2.2.3. One day before the experiment, aspirate 14 female mosquitoes per acrylic plate and anesthetize them at 4 °C. Carefully place one mosquito at a time into each of the wells with a feather or aspirator.
    2.2.4. Tape the side of the acrylic plate with Scotch tape to hold the lid in place. Do not fold the tape over the top or bottom of the acrylic plate, as it would obstruct the view of the camera.
    2.2.5. Place the acrylic plate on top of three cotton wicks soaked with 400µM ATR in distilled water (See step 1.5) placed in parallel. Align each of the 3 rows to a wick so all the mosquitoes have access to the water-ATR solution. Place the acrylic plate in a dark incubator overnight.
  2.3. **Running the trial**
    2.3.1. Connect the camera, LEDs, and Arduino to the computer. Turn on the power supply. Wipe down the surface of the opto-thermocycler with 70% ethanol and allow it to dry.
    2.3.2. Using a dark container, transfer the acrylic plate of mosquitoes to the middle of the PCR block. Cover the opto-thermocycler with a blackout curtain.
    2.3.3. Open SpinView on the computer. Open the camera view under the **Cameras** tab. Plug in the infrared lights and make sure the PCR block and acrylic plate are visible in the camera view. If needed, adjust the acrylic plate so it is aligned, centered, and the thermocouple is visible in the bottom right corner of the acrylic plate. Ensure the synchronization light is in frame (**Figure 2C**).
    2.3.4. Adjust the settings on the camera and in SpinView to ensure a crisp, cropped video showing the acrylic plate. In the **Image Format** tab, adjust the **Width, Height, Offset X**, and **Offset Y** options to ensure the acrylic plate and synchronization light are visible. In the **Settings** tab, set **Acquisition Frame Rate Enable** to true and **Acquisition Frame Rate** to 30 fps. Turn **Exposure Auto** and **Gain Auto** off. Adjust the **Exposure Time** and **Gain** values to create a clean contrast between the black background and the mosquitoes.
    2.3.5. Click the red record button on the upper right side which opens a separate window to save the video file.
    2.3.6. In the pop-up window, set the file path at the top: Click **Browse** > Navigate to the experiment folder > Create a new folder with the trial name with the appropriate folder name.
    2.3.7. Set the number of frames to record to the appropriate number for the experiment (e.g., 43500) or to 0 to manually stop the recording.
    2.3.8. In the **Videos** tab, set the following fields: **Video Recording Type**: Mjpg, **Video File Split Size**: 1000 MB, **JPEG Compression Quality**: 65.
    2.3.9. Open the ‘optothermo_lightheat_random1’ program on the thermocycler and start it. The thermocycler should begin in a hold step at 25 °C.
    2.3.10. Open the matching program, ‘TC_optothermo_lightheat_random1.ino’, on the Arduino IDE, and upload it to the board. Open the serial monitor and ensure that the temperature is approximately 25 °C.
    2.3.11. Close the serial monitor in the Arduino IDE. Open CoolTerm to begin the acquisition of the Arduino serial monitor output. Make sure the serial port and channel match the serial port and channel on the Arduino IDE. The output should start showing on the CoolTerm window.
    2.3.12. Begin the recording of the Arduino output to a Text/Binary file. Start the video recording on Spinview. Click **Resume** to end the hold on the thermocycler. Watch the Arduino output on CoolTerm to check if the temperature dip to 18 °C. was detected and the stimulus counter increased to 1, initiating the trial. Allow the assay to run until the trial is completed.
  2.4. **Clean up**
    2.4.1. Stop the video recording and CoolTerm output acquisition. Close out of both programs. End the thermocycler program. **NOTE**: This is recommended even when running another trial immediately after to clear cache.
    2.4.2. Take the acrylic plate of mosquitoes from the thermocycler and place them in a −20 °C freezer overnight to euthanize them.
    2.4.3. If starting another trial, repeat section 2.3.
    2.4.4. Unplug the camera and Arduino cables from the computer. Turn off the power supply. Unplug the infrared LEDs from the outlet.
    2.4.5. Clean acrylic plates by removing the Scotch tape and discarding the dead mosquitoes in biohazard waste. Spray the lid and acrylic plate with 70% ethanol and wipe all the sides of the wells and mesh with a gloved finger carefully. Rinse thoroughly with deionized water and stack acrylic plates for storage with a paper towel separating lids to avoid scratching.
3. **Blood Blanket Assay**
  3.1. **Assembling components of the assay**
    3.1.1. Construct the opto-thermocycler assay as described in section 2.1. The blood blanket assay will use the components of the opto-thermocycler with some modifications. Refer to the figure for a diagram of the assay (**Figure 3A**), thermal stimulus (**Figure 3B**), and aluminum plate (**Figure 3C**).
    3.1.2. Construct an aluminum blood meal feeding plate as seen in **Figure 3C**.
      3.1.2.1. Using a laser cutter or waterjet cutter, construct the top and bottom of the plate. Construct the bottom of the plate by cutting an aluminum sheet 0.8 mm (1/32 inch) thick to 90 mm by 134 mm. Construct the top by cutting an aluminum sheet 1.6 mm (1/16 inch) thick to 90 mm by 134 mm. Cut 15 rectangular holes (17.2 mm by 18.6 mm) into the top plate. The template files can be found in the blood blanket GitHub repository (https://github.com/sorrellslab/bloodblanket) under “Aluminum plate templates”.
      3.1.2.2. Sandwich between the aluminum parts a thin silicone gasket (0.5 mm thick) to prevent leakage. Cut using a laser cutter in the same shape as the aluminum top.
      3.1.2.3. Attach layers together by eight 0.47625 cm (3/16 inch), dome-topped hex 4-40 standard bolts. Use a forming tap to create threads for the bolts in the metal pieces. Connect these through round and “W” washers on the bottom of the plate for added security when placed on top of the thermocycler block.
    3.1.3. Download the Arduino code and thermocycler program for the blood blanket assay from the GitHub repository (https://github.com/sorrellslab/bloodblanket).
    3.1.4. Prepare stock solutions.
      3.1.4.1. Prepare aliquots of 20 mM ATP in 25 mM NaHCO_3_ and store at −20 ° C until use. **NOTE**: Prepare a fresh ATP solution every six months. Avoid repeated freeze-thaw cycles.
      3.1.4.2. Prepare 1 M NaHCO_3_ and 5 M NaCl (store each at room temperature). **NOTE**: NaHCO_3_ is only stable in solution for 7 days.
  3.2. **Experiment Preparation**
    3.2.1. Rear mosquitoes and transfer to acrylic plates as described in sections 1 and 2.2. If comparisons are going to be made between different genotypes, blind them to the experimenter.
    3.2.2. The day of the experiment, prepare a 10 mL minimal artificial blood meal (110 mM NaCl, 20 mM NaHCO_3_, and 1.5 mM ATP) by adding 750 µL of 20 mM ATP, 220 µL of 5 M NaCl, and 200 µL of 1 M NaHCO_3_ stock solutions to 8.83 mL of deionized water. Mix thoroughly by inverting. **NOTE**: Make the artificial blood meal fresh prior to each experiment. Minimal blood meal is used for ease of clean up..
    3.2.3. Set up SpinView recording according to steps 2.3.4 to 2.3.8.
    3.2.4. Open “Red-light-Only BB_.ino” Arduino file (or desired stimulus file) for the experiment.
  3.3. **Running Trials**
    3.3.1. Pipette 650 µL of the minimal artificial blood meal to each well of the plate, ensuring that the meal entirely fills each well with no overflow.
    3.3.2. Stretch a 4 inch x 4 inch square of parafilm over an 8 inch diameter ring. Place stretched parafilm over the plate, pressing on the edges until the parafilm sticks to the plate. Cut with a scalpel about 0.5 inch around the plate and tuck the excess parafilm underneath the plate to seal in the meal.
    3.3.3. Place the plate directly on top of the PCR block to allow maximum heat transfer. Ensure that the plate is firmly in contact with the block as shown in **Figure 3A**.
    3.3.4. **NOTE**: Blackout tape is used on the surface of the PCR block for the opto-thermocycler but not for the blood blanket.
    3.3.5. Place the thermocouple onto the surface of the parafilm in the bottom right corner of the aluminum plate. Connect the camera, LEDs, and Arduino to the computer. Turn on the power supply.
    3.3.6. Start the “BB_thermocycler_program.x50prog” program on the thermocycler. **NOTE**: The rate of temperature changes is the maximum ramp speed for the thermocycler, chosen to best mimic landing of mosquito onto host for feeding.
    3.3.7. Using a dark container and dim room lights to minimize light exposure, transfer the acrylic plate of mosquitoes onto the aluminum plate, ensuring each well is aligned with the mosquitoes. Ensure the thermocouple is in the bottom right corner of the plate and is in contact with the parafilm to measure the temperature of the artificial blood meal.
    3.3.8. Start the program by re-uploading the “Red-light-Only BB_.ino” program in the Arduino IDE. Close the serial monitor. In CoolTerm, press **Connect** to see the current temperature. Using this starting temperature, adjust the trigger temperature in the Arduino script to be approximately 0.6 °C below the starting temperature.
    3.3.9. In the CoolTerm window, select **Capture to Text/Binary file** and start data collection.
    3.3.10. Start video recording in the record window in SpinView.
    3.3.11. Click **Resume** to end the hold on the thermocycler. Watch the Arduino output on CoolTerm to ensure that the heat dip was detected and the trial has started.
    3.3.12. After the duration of the trial, click **Stop Recording**. On CoolTerm in the **Capture to Text/Binary file** tab, select **Stop** to writing the serial output.
    3.3.13. Remove the plate of mosquitoes from the blood blanket plate and move them to the cold room to anesthetize them. Count the number of mosquitoes with enlarged abdomens that contain the artificial blood meal as engorged, including those who have partially fed.
    3.3.14. Between trials, clean the surface of the PCR block by wiping with a Kimwipe moistened with 70% ethanol. Remove the parafilm and rinse the aluminum plate with deionized water.
    3.3.15. Repeat section 3.3 for further trials. Rotate the order of trials between days to minimize the effect of circadian activity cycles.
  3.4. **Clean up**
    3.4.1. Stop recording in all programs and quit. Unplug USB cords for the camera, Arduino, and infrared lights. Shut down the computer. Stop the “thermocycler_program.x50prog” program running on the thermocycler.
    3.4.2. Place mosquito plates in a −20 °C freezer overnight to kill the mosquitoes.
    3.4.3. Wash the aluminum plate with deionized water. Douse the acrylic plate with 70% ethanol, removing any debris and residue with gloved fingers from the wells, then rinse with deionized water and leave to dry overnight before loading more mosquitoes. **NOTE**: Plates should be dried overnight after washing to make sure there is no moisture left before loading mosquitoes. If there is moisture, this will cause condensation on the lid and will interfere with the video.
4. **Opto-membrane Feeder Assay**
  4.1. **Assembling components of the assay**
    4.1.1. Assemble a frame using opto-mechanical components and a 30.48 cm (12 inch) x 30.48 cm (12 inch), black, 0.635 cm (1/4 inch) thick, acrylic platform. Create a hole in the center of the acrylic with a diameter of 11.43 cm (4.5 inch) using a laser cutter. Cut holes 2.54 cm (1 inch) from each of the corners, to allow the acrylic to rest on the supports of the frame, approximately 23.495 cm (9.25 inch) above the base. In the center, attach a black acrylic ring with a 19.05 cm (7.5 inch) inner diameter and a 20.32 cm (8 inch) outer diameter (**Figure 4A**). Templates for all acrylic components are in the opto-membrane feeder GitHub repository (https://github.com/sorrellslab/opto-membranefeeder/releases/tag/v1.0.0) along with the Arduino programs. See figure for overall setup (**Figure 4A**), stimulus (**Figure 4B**), circuit diagram (**Figure 4C**), and video still image (**Figure 4D**). **NOTE**: The specific materials used to assemble this platform are not critical, so more cost-effective options may be available.
    4.1.2. Attach a camera with an infrared long-pass filter below the center hole, pointing up toward the mosquito canister. Adjust the aperture to be fully open, allowing selective focus on the mesh top of the canister (**Figure 4A**).
    4.1.3. Construct canisters to contain the mosquitoes (**Figure 4A**).
      4.1.3.1. Cut a clear, polycarbonate tube with a diameter of 11.43 cm (4.5 inch) into 12.7 cm (5 inch) cylindrical segments.
      4.1.3.2. Build bottoms for the canisters by laser-cutting circles with a 11.43 cm (4.5 inch) diameter from 0.3175 cm (1/8 inch) thick acrylic. The edges of the circles are etched with the laser so that they are inset into the tube. Attach to the tubes with plastic epoxy.
      4.1.3.3. Create an inset lid using black 0.635 cm (1/4 inch) and 0.3175 cm (1/8 inch) acrylic and UV-resistant black mesh. The inner ring has a diameter of 10.795 cm (4.25 inch), while the outer ring has a diameter of 12.065 cm (4.75 inch), with the black mesh stretched over the smaller ring. The mesh should be sewn on through laser cut holes with waxed thread. **NOTE**: Canisters should always be placed on paper towels to avoid scratching the acrylic bottoms through which video is recorded.
    4.1.4. Construct acrylic containers for blood meal delivery (**Figure 4A**).
      4.1.4.1. Cut a ring with a 5.08 cm (2 inch) inner diameter and 6.6 cm (2.6 inch) outer diameter from 0.15875 cm (1/16 inch) clear acrylic.
      4.1.4.2. Attach this to another clear ring with a 5.842 cm (2.3 inch) inner diameter and 6.604 cm (2.6 inch) outer diameter using acrylic glue.
    4.1.5. Place a metal mesh cylinder with a 13.97 cm (5.5 inch) height and a 19.05 cm (7.5 inch) diameter inside the black acrylic ring at the center of the platform.
    4.1.6. Inside the metal mesh, attach a coil of RGB LED lights spaced 3.81 cm (1.5 inch) from the exterior of the canister, to be controlled by an Arduino Uno board which rests on the black acrylic platform (**Figure 4C**). Using a power meter, measure light intensity throughout the inside of the cylinder and adjust the location of the lights so the arena is illuminated between 3.5-6 µW/mm^2^.
    4.1.7. Outside the metal mesh, attach a ring of 850 nm infrared LEDs to the frame using zip ties, facing inward.
    4.1.8. Plug a USB cord into the Arduino Uno board and connect to the computer that will be used to run the software. Connect a USB cord from the camera to the computer. Turn the off the connection to the internet off on the computer.
    4.1.9. Place the opto-membrane structure in a dark incubator or environmental room at 26 °°C and 80% relative humidity for the duration of the experiments.
  4.2. **Preparing the mosquito canisters**
    4.2.1. Prior to the experiment, wash mosquito canisters by spraying 70% ethanol and wiping gently with a soft sponge. Rinse the canisters with deionized water and air-dry overnight.
    4.2.2. Two days prior to the experiment, sex mosquitoes under cold anesthesia and place 20 females into each canister. Provide access to ATR according to step 1.5.
    4.2.3. At this time, genotypes should be blinded to the experimenter. The order of the trials should be rotated between days to minimize differences in behavior outcomes due to experimental timing.
  4.3. **Running Trials**
    4.3.1. Prepare the blood meal.
      4.3.1.1. Place 119 mL (4 oz) bottles of water into a 45 °C heat bath, one for each trial. This volume and temperature maintain the blood meal close to body temperature for the duration of the experiment.
      4.3.1.2. Prepare 5 mL aliquots of defibrinated sheep blood. Store at 4 °C and invert prior to use to prevent separation. **NOTE**: Defibrinated sheep blood is used in the blood meal. Make the blood meal fresh prior to each experiment. It should be handled while wearing gloves and a lab coat. Dispose of single use materials in a biohazard bin. All reusable materials that contacted blood should be soaked in a 10% bleach solution and rinsed with DI water.
      4.3.1.3. Place first blood aliquot into a 45 °C heat bath for at least 20 minutes prior to the trial.
      4.3.1.4. Thaw ATP on ice.
      4.3.1.5. Stretch a 5.08 cm (2 inch) x 5.08 cm (2 inch) square of parafilm over the acrylic lid on the side with the larger, flat circle, creating a well for the blood meal (see **Figure 4A**).
    4.3.2. Wipe the computer with 70% EtOH to remove human odors. Turn on the computer and connect it to an external hard drive. Connect the camera and the Arduino to the computer. Plug the RGB and infrared lights into a power source.
    4.3.3. Open the SpinView software and the Arduino file for the appropriate stimulus.
    4.3.4. Create a folder on the external hard drive to contain the experiment data.
    4.3.5. Upload the “blue_only.io” Arduino file to provide dim blue 471 nm light from the RGB LEDs to the arena (following **Figure 4B**).
    4.3.6. Without exposing mosquitoes to excess CO_2_ from breath, examine the canister for the trial and record any deaths prior to the experiment.
    4.3.7. Place the canister on the center of the platform and allow acclimation under dim blue light for 10 to 20 minutes prior to stimulus exposure. **NOTE**: The acclimation step and experiment are run under dim blue light because complete darkness inhibits mosquito flight.
    4.3.8. Enable video capture.
      4.3.8.1. In the SpinView program, select the camera. In the **Settings** tab, adjust the exposure and gain as needed to maintain high contrast between the light-colored insects and the dark background (**Figure 4D**).
      4.3.8.2. Click the red record button on the top bar to adjust video capture settings.
      4.3.8.3. Set the file path by clicking **Browse** and navigating to the external hard drive.
      4.3.8.4. Create a new folder with a descriptive name for each trial. Do not include spaces in the folder name.
      4.3.8.5. In the **Save Options** tab, select **Capture 0 frames**, which will allow video to record until manually stopped. Under **Recording Mode**, select **Buffered**. In the **Videos** tab, set the following fields: **Video Recording Type**: Mjpg, **Video File Split Size**: 1000 MB, and **JPEG Compression Quality**: 65.
    4.3.9. Immediately before stimulus exposure, pipette 500 µL of 20 mM ATP, for a final concentration of 2 mM ATP, into the prepared 5 mL blood aliquot. Invert the tube several times to mix.
    4.3.10. Pour the ATP-blood solution into the blood meal container.
    4.3.11. On top of the blood meal, place an inverted 119 mL (4 oz) bottle filled with 45 °C water to maintain the temperature of the blood close to human body temperature throughout the experiment.
    4.3.12. In the SpinView software, select **Start Recording**.
    4.3.13. Without stimulating mosquitoes with breath, place the blood meal on top of the mesh lid of the canister so mosquitoes can pierce the parafilm membrane through the mesh.
    4.3.14. Upload the Arduino program for “1sON_10sOFF.io” (**Figure 4B**), or the desired stimulus. This will deliver 1 second of 624 nm red light every 10 seconds, until the “blue_only.io” file is manually uploaded. The wavelength of 624 nm is sufficient to opto-genetically activate CsChrimson.
    4.3.15. Place the blood aliquot into the heat bath for the following trial.
    4.3.16. After the trial has run for 15 minutes, select **Stop Recording**.
    4.3.17. Upload the “blue_only.io” file to stop the red-light stimulus.
    4.3.18. Remove the blood meal holder and water bottle. Remove the canister and anesthetize the mosquitoes at 4 °C.
    4.3.19. To perform additional trials, return to section 4.3.
    4.3.20. Visually examine the anesthetized mosquitoes to determine and record the number that engorged on blood by looking for a blood-filled abdomen.
  4.4. **Clean up**
    4.4.1. Close all programs.
    4.4.2. Eject external hard drive and disconnect from the computer.
    4.4.3. Shut down the computer.
    4.4.4. Unplug all cords connected to the computer and power sources.
    4.4.5. Clean the mosquito canisters as previously described in step 4.2.1.
    4.4.6. Clean acrylic blood meal holders
      4.4.6.1. Fill a half-sized polycarbonate pan 2/3 full with water and add 100 mL of bleach.
      4.4.6.2. Add acrylic blood meal holders and plastic bottles to the pan. Cover with a lid and let soak for 10 minutes.
      4.4.6.3. Rinse bleach and remove parafilm. Scrub gently and rinse to remove any blood remnants.
5. **Analysis of behavior**
  5.1. Prepare the videos for analysis by compressing them into .mp4 files using ffmpeg (version 8.0.1).
  5.2. If the video files are split, merge them into one video file using any video editing software.
  5.3. Open the training videos in SLEAP (version 1.4.1) in grayscale. If the video recording settings are appropriate, everything should be within frame and moving mosquitoes should not be blurry. **NOTE**: Other body part tracking software can be used. The SLEAP version used in this paper is 1.4.1. Refer to SLEAP documentation for up-to-date protocols.
  5.4. Create a skeleton that labels the relevant parts of the animals according to the behaviors being assessed (**Figure 2D, 3D, and 4E**).
  5.5. Using the “Random frame” option under the **Suggested frame** tab, randomly pick 10 frames and label each mosquito as an instance. If certain parts of the mosquito are obstructed (i.e. the proboscis tip is inserted into the mesh during probing), label your best guess for the location of the body part. Train a model on the labeled frames with these settings:
    5.5.1. For a Top-Down multi-animal model (Used for the opto-thermocycler and blood blanket model): In the **Training Pipeline** tab, set **Max Instances** to 14. In the **Centroid Model Configuration** tab, set **Crop Size** to **Auto**, set **Plateau Min. Delta** to 1e-04, set **Plateau Patience** to 10, set **Stride** to 8, and the **Anchor Part** as thorax.
    5.5.2. For a Bottom-Up multi-animal model (Used for the Opto-membrane feeder): In **the Training Pipeline** tab, set **Max Instances** to no max. In the **Bottom-Up Model Configuration** tab, set **Crop Size** to **Auto**, set **Plateau Min. Delta** to 1e-04, set **Plateau Patience** to 10, and set **Stride** to 16.
    5.5.3. Predict on a set number of frames with these settings: In the **Inference Pipeline** tab, for the opto-thermocycler and blood blanket, set **Max Instances** to 14, **Tracker** to simple, **Max number of tracks** to 14, and **Connect Single Track Breaks** to **True**. For the opto-membrane feeder, set **No Max** for **Max Instances** and **Max number of tracks, Tracker** to simple, and **Connect Single Track Breaks** to True.
  5.6. Correct the predicted frames and re-train. Iterate through cycles of prediction, labeling, and training on the training data set. Validation was done by spot-checking frames by hand at random and making note of consistent errors (i.e. predicting a mosquito where no mosquito is present, predicting a single mosquito twice, predicting the wrong orientation, incorrect body parts, etc.) Focus on frames where these errors occur. This can take up to 150 labeled frames, depending on the number of edge cases in your videos. SLEAP also provides evaluation metrics for its models within the GUI, which provides another criterion for prediction validation. The SLEAP models used in this study are made available as a resource on each assay’s respective GitHub repositories Use the model on experimental videos once it is able to predict at a high enough accuracy and precision, based on visual inspection, and export into .csv or .h5 for behavior-tracking analysis.
  5.7. Behavior classifiers can subsequently be trained using pose estimation outputs as described previously^8^. Behaviors are defined as follows. Walking is translational body movement along with leg motion. Flying is rapid translational body movement with wings extended. Grooming is repetitive movement of legs rubbing against body parts including antennae, proboscis, wings, and other legs. Probing is insertion of the proboscis through mesh at the bottom of the container. Behavior classifiers should be accurate at >90% when compared to human annotations.
  5.8. Perform statistical analysis by choosing non-parametric tests as the data are not typically normally distributed. Perform the corresponding post-hoc tests with correction for multiple comparisons if the null hypothesis is rejected at a significance threshold of ***P*** < 0.05. For engorgement data, experimental replicates are groups of mosquitoes, whereas for behavior data, replicates are individual mosquitoes.

**Figure 2:**
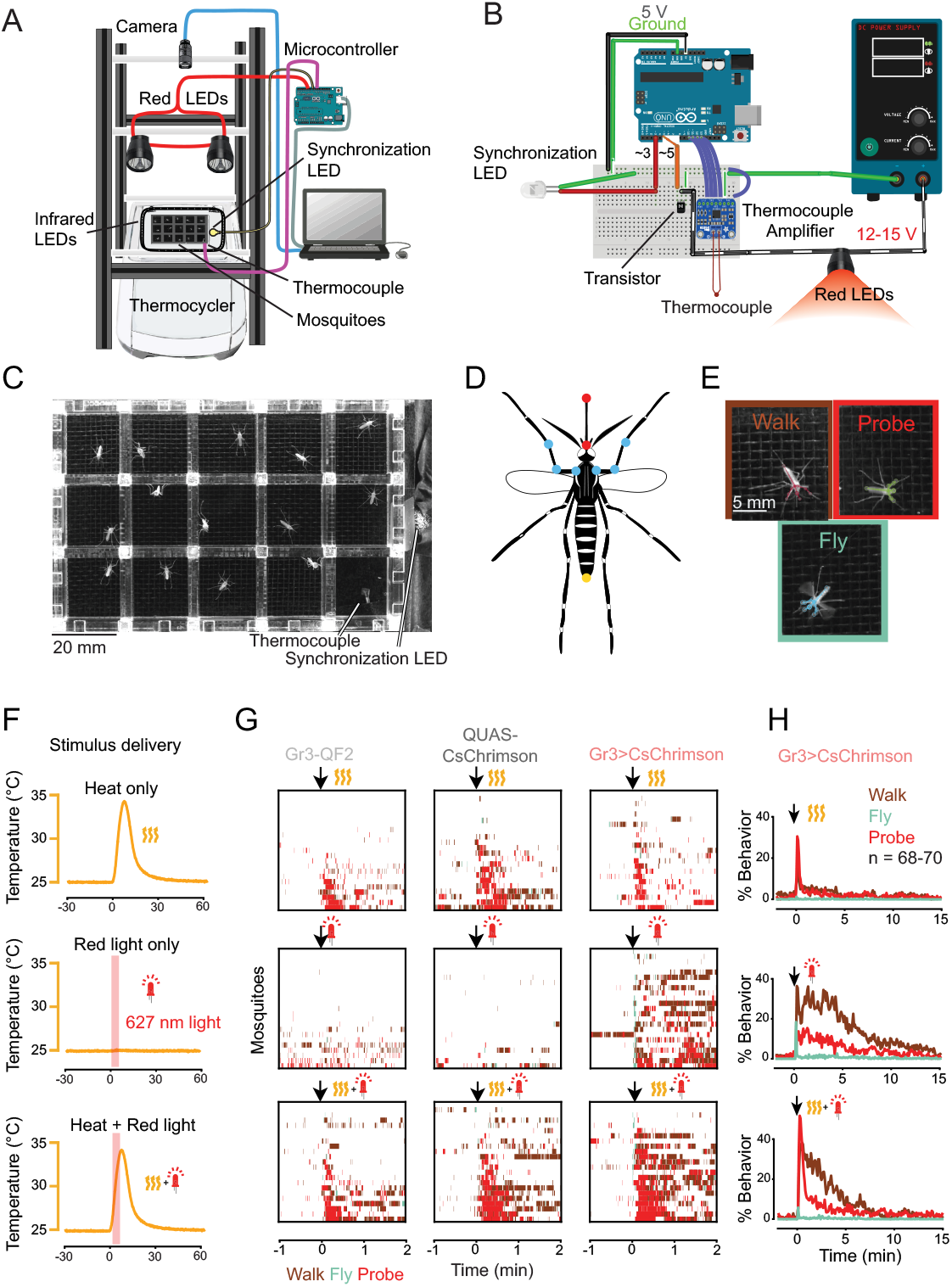
The opto-thermocycler quantifies arousal and search behaviors during optogenetic activation. **(A)** Schematic of opto-thermocycler assay consisting of a plate of mosquitoes on the block of a PCR thermocycler with overhead LEDs and a camera. **(B)** Circuit diagram for the microcontroller that detects the thermocycler temperature and controls the LEDs. **(C)** Still image from the assay depicting the acrylic plate that holds up to 14 mosquitoes. The thermocouple is in the bottom-right well. **(D)** Mosquito skeleton used in SLEAP model to estimate poses. Points labeled on the body include: the tip and base of the proboscis, used to detect probing behavior (shown in red); the front legs where they connect to the thorax, where the femur joins the tibia and where the tibia joins the tarsus (shown in blue); and the tip of the abdomen (shown in yellow). A top-down multi-animal model was used. Pose estimators and behavioral classifiers were used to quantify mosquito arousal and probing behavior. **(E)** Examples of mosquito walking, flying, and probing, with SLEAP predictions overlaid on top. Walking and flying are used as a behavioral metric for arousal. **(F)** Output of stimulus delivery. Yellow wavy lines indicate body temperature heat stimuli and red LEDs indicate red light stimuli. **(G)** Ethograms of walking, flying, and probing behavior by Gr3-QF2, QUAS-CsChrimson, and Gr3>CsChrimson mosquitoes (n = 22-23 individual mosquitoes per genotype) following stimulus delivery. **(H)** Long-lasting behavior following optogenetic activation in Gr3>CsChrimson mosquitoes (n = 68-70 individual mosquitoes). Data in **(F-H)** are reproduced from a previous publication^8^.

**Figure 3:**
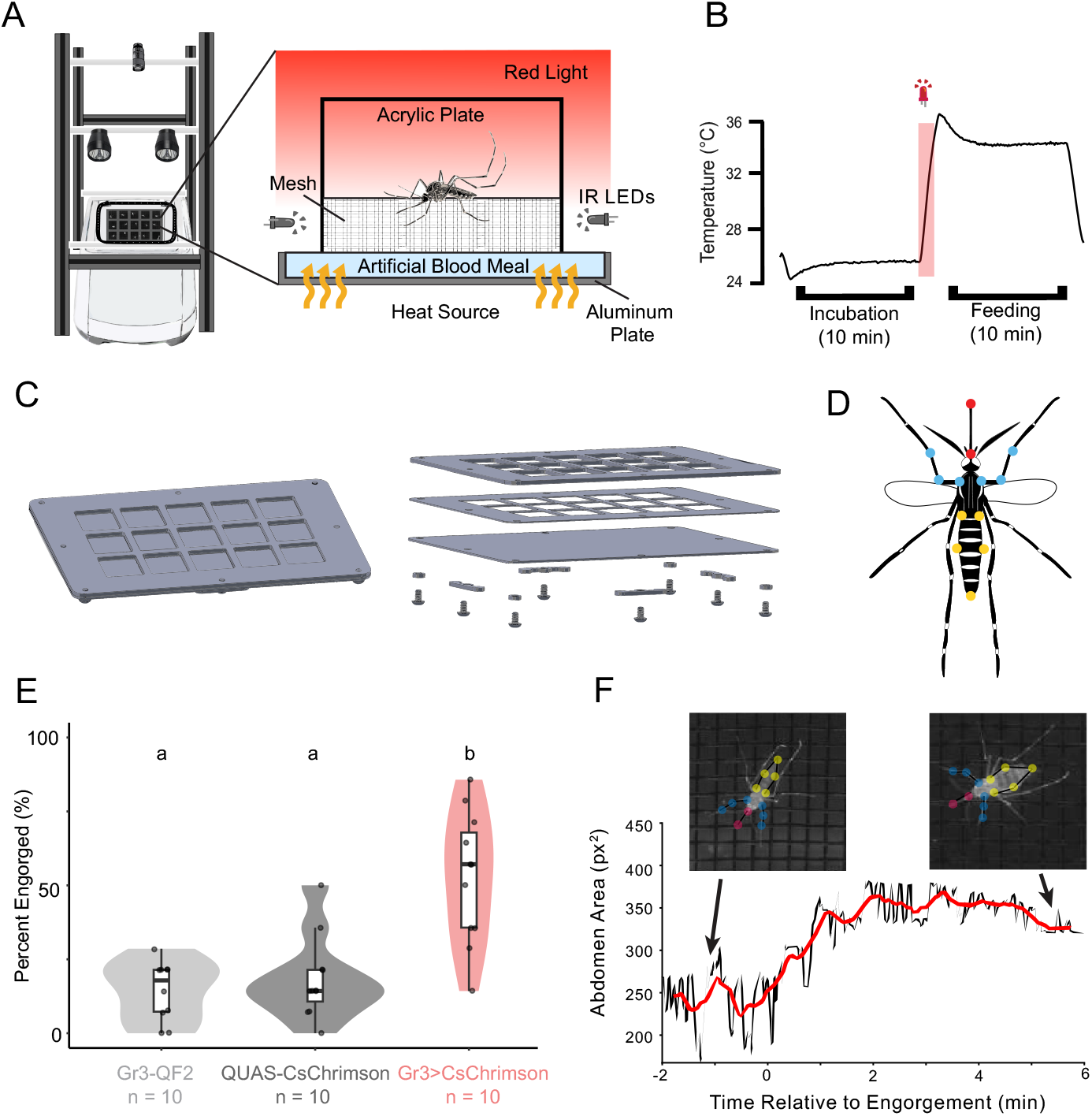
The blood blanket assay quantifies engorgement to optogenetic activation. **(A**) Schematic of the blood blanket assay. The mosquito is contained in an acrylic cage where accessible artificial blood is presented beneath a mesh screen. **(B**) Readout of temperature and red light stimulus from the assay. **(C**) Schematic of aluminum blood blanket feeding plate assembly. **(D**) The SLEAP skeleton used for pose estimation included the points described in **Figure 2D** in addition to four points along the sides of the abdomen where they meet the thorax and the edges of the thickest part of the abdomen. Engorgement was measured as the area within the five points along the abdomen perimeter using predictions from a top-down multi-animal model. **(E**) Violin plot depicting the mean percent engorged on the artificial blood meal per genotype (n = 10-11 trials of 14 mosquitoes per trial for each genotype). Significance was determined using the Kruskal-Wallis test, followed by a Dunn Test with Holm correction. Different letters indicate a significance of p < 0.05 between groups. **(F**) Engorgement over time of a Gr3>CsChrimson animal as measured by abdomen area. The red trendline displays a 30-second sliding window over eight minutes of the trial, with x = 0 marking time when engorgement begins.

**Figure 4:**
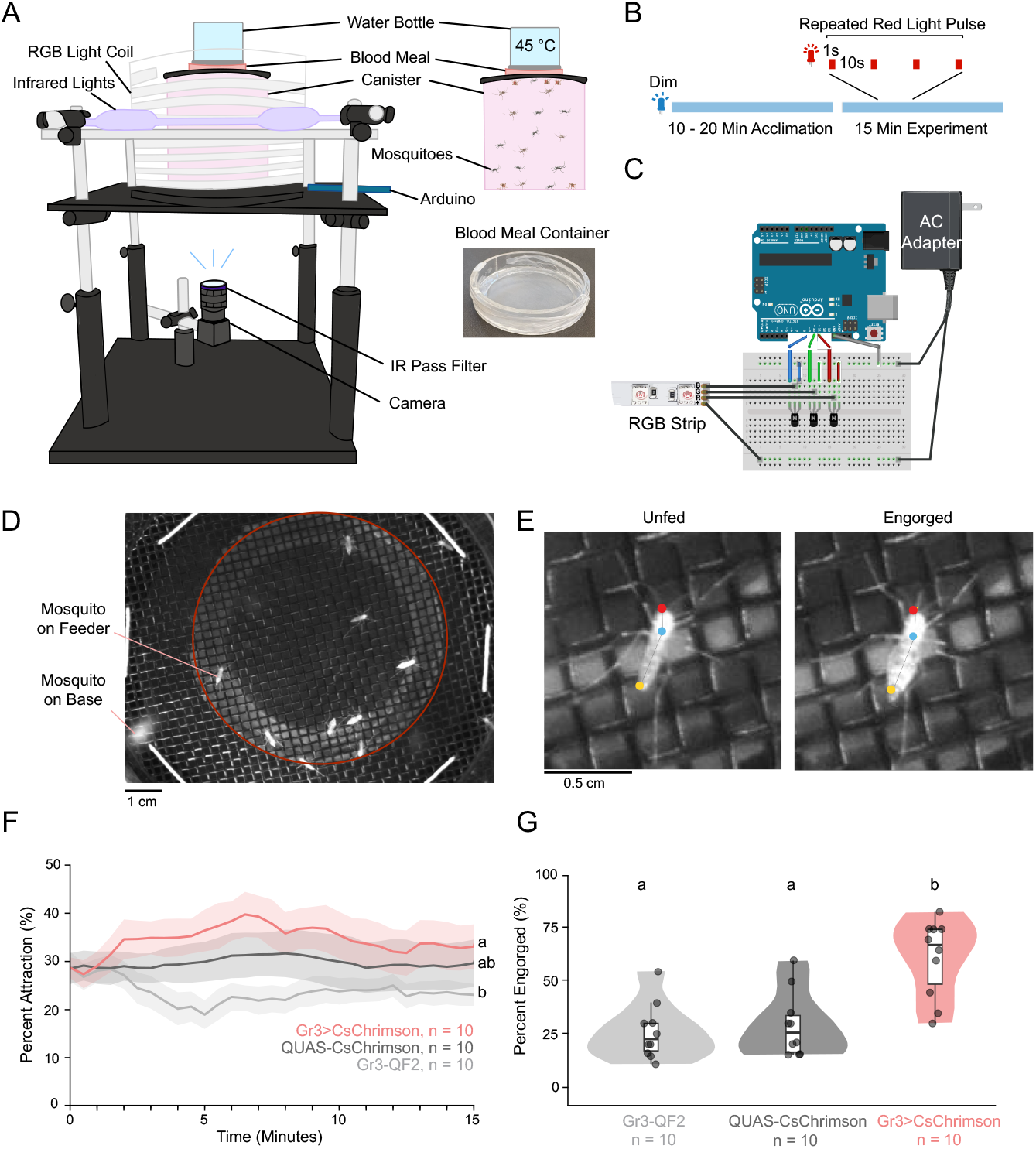
The opto-membrane feeder assay measures attraction and blood feeding. **(A)** Schematic of the opto-membrane feeder with individual components. **(B)** Timeline of experiment trials. The blue line represents dim blue light throughout the acclimation period and experiment. A segment of the experimental stimulus is shown, representing 1-second pulses of red light spaced 10 seconds apart. **(C)** Diagram depicting the wiring for the opto-membrane feeder. **(D)** Example frame from the opto-membrane feeder video. Mosquitoes in the red circle are counted to determine percent attraction. **(E)** Examples of an unfed mosquito and an engorged mosquito, with the head, thorax, and abdomen labeled by the SLEAP tracking model for the opto-membrane feeder. A bottom-up multi-animal SLEAP model was used to measure attraction to the region of interest where the blood meal is placed. **(F)** Line graph depicting mean percent occupancy on the blood meal region (which is used as a behavioral metric for landing) over time, by genotype (n = 10 trials of 20 mosquitoes per genotype). Mean is plotted every 30 seconds. Shaded regions depict standard error. **(G)** Violin plot depicting the mean percent engorged on a blood meal by genotype. In **(F, G)**, Significance determined by the Kruskal-Wallis test, followed by the Dunn Test with Holm correction. Different letters indicate significant differences between groups.

## Results

Three behavior assays were used to observe host-seeking and blood-feeding behaviors in mosquitoes as shown in **Figure 1A**. Mosquitoes were reared under blue light-dark cycle (14:10 LD; **Figure 1B**) to synchronize circadian rhythms so experiments could be run during wake periods. Two days before experiments, mosquitoes were transferred to total darkness and provided with the opsin cofactor ATR. If mosquitoes are asynchronous or not fed ATR, they will show low activity in behavioral assays. The genotypes used for each assay were Gr3>CsChrimson and two genetic controls: the driver line Gr3-QF2 and the effector line QUAS-CsChrimson.

The opto-thermocycler assay (Section 2) allows for the assessment of mosquito arousal and probing behavior over long periods of time in response to heat and light stimuli. Heat delivery mimics the mosquito approaching a warm-blooded host while the 5-second light delivery mimics a long exhalation of breath^33^. The use of a thermocycler allows for quick surface heating and cooling that is synchronized with optogenetic activation via a thermocouple and microcontroller (**Figure 2A, B**). The three mosquito genotypes were placed in acrylic plates with isolated wells with mesh bottoms in which they demon strate walking, flying, and probing behavior (**Figure 2C**). The position of mosquito body parts was detected using pose-tracking algorithms, such as SLEAP^34^ and DeepLabCUT^35^ (**Figure 2D, E**). Behavior classification algorithms can be used starting from pose tracking (e.g. SimBA^36^, A-SOiD^37^) or video (e.g. FERAL^38^) input. **Figure 2E** shows the SLEAP labels during flying, walking, and probing, essential host-seeking and feeding behaviors. These behaviors can be classified by the velocity of the thorax and the distance between the base and tip of the proboscis, respectively. Here, we show the output of behavior tracking using APT and JAABA^39^. Mosquitoes were exposed to either a heat increase, a light stimulus, or both heat and light stimuli simultaneously, then a break of 20 minutes before the next stimulus^8^. (**Figure 2F**). When exposed to heat, all three genotypes exhibit a short period of probing (**Figure 2G**). Importantly, when exposed to a 5-second light stimulus, the Gr3>CsChrimson mosquitoes become aroused or “activated,” showing increased walking and flying behavior that lasts up to 10-15 minutes. Mosquitoes also exhibit probing, which is defined as the appearance and disappearance of the proboscis into the mesh bottom of the plates (**Figure 2G, 2H**). The opto-thermocycler allows for repeated, precise optogenetic and heat stimulation to multiple individuals, making it a useful assay to study arousal and probing behavior.

The blood blanket assay (Section 3) is built upon the opto-thermocycler assay but provides a palatable artificial minimal blood meal so mosquito engorgement can be quantified (**Figure 3A**). A 5-second pulse of red light (~627 nm) was administered to optogenetically activate CO_2_ sensory neurons and the thin layer of artificial blood meal was heated from 25 °C to ~35 °C (as measured at the surface by a thermocouple; **Figure 3B**). The three genotypes were each presented with an artificial blood meal in an aluminum feeding plate (**Figure 3C**). Gr3-QF2, QUAS-CsChrimson, and Gr3>CsChrimson mosquitoes (n = 10-11 trials per genetic group, 14 mosquitoes each) were each presented with an artificial blood meal in an aluminum feeding plate (**Figure 3C**). To measure engorgement of the mosquitoes over time throughout the experiment, SLEAP was used to track points on the body of the mosquito. This was done by creating a skeleton that included 5 nodes on the abdomen of the mosquito (**Figure 3D**). At the end of each trial, the number of mosquitoes engorged was manually scored. The control mosquito lines, Gr3-QF2 and QUAS-CsChrimson, demonstrated a similar average engorgement (**Figure 3E**, *P* = .401). In contrast, the Gr3>CsChrimson line demonstrated significantly higher rates of feeding, with an average engorgement rate of 53% (**Figure 3E**, *P* = .00078 versus Gr3-QF2 and *P* = .00097 versus QUAS-CsChrimson). Individual animal engorgement over time was identified as the change in the abdominal area enclosed by 5 nodes (**Figure 3F**). These results show that the blood blanket assay can be used to assess the effects of optogenetic activation of CO_2_-responsive neuronal populations on the engorgement behavior of mosquitoes on an artificial blood meal.

The opto-membrane feeder assay (Section 4), where mosquitoes are presented with access to a warm blood meal in a cylindrical arena, allows for study of mosquito attraction to a blood source and engorgement. To determine if optogenetic activation of CO_2_ sensory neurons is sufficient to cause attraction to and feeding on a blood source, Gr3>CsChrimson and control mosquitoes (n = 10 for each genotype, where n refers to the number of trials run, with 20 mosquitoes per canister per trial) were presented with access to a warm blood meal in the opto-membrane feeder under exposure to a repeated 1-second pulse of bright red light every 10 seconds for a duration of 15 minutes (**Figure 4B**). To measure attraction to the membrane feeder, a SLEAP model with three points was used to track the head, thorax, and abdomen of landed mosquitoes, counting those within the region of interest (**Figure 4D, Figure 4E**). Gr3>CsChrimson mosquitoes showed increased attraction to the blood meal region over time compared to the Gr3-QF2 control (**Figure 4F**, *P* = .027). The Gr3>CsChrimson mosquitoes did not show statistically higher attraction than the QUAS-CsChrimson (**Figure 4F**, *P* = .310). The control genotypes Gr3-QF2 and QUAS-CsChrimson showed similar engorgement of 26% and 29% respectively, manually scored as described in step 4.3.20 (**Figure 4G**, *P* = .351). In contrast, Gr3>CsChrimson mosquitoes demonstrated a significantly higher percentage of engorgement than either control, at an average rate of 61% (**Figure 4G**, *P* < .001 versus Gr3-QF2 and *P* = .002 versus QUAS-CsChrimson). These results indicate that the opto-membrane feeder assay is a viable means to assess the effects of optogenetic activation of CO_2_ neurons on mosquito attraction and engorgement on a blood meal.

## Discussion

Here, we have described detailed protocols for assaying the sufficiency of neuronal cell types to drive steps in host attraction and biting. Biting behavior consists of a series of behavioral steps from long-range detection to short-range attraction, piercing of skin, and engorgement. In principle, neuronal types could control specific actions or steps, bias behavior toward specific sequences, increase the duration of the whole behavior, or play many other roles. Thus, the multiple assays we present are necessary to dissect the function of neurons in this process.

Optogenetic tools have been created that are sensitive to wavelengths from blue (~450 nm) to red (~650 nm), spanning the visible spectrum. Red light is particularly useful in insects as they typically have low sensitivity and/or behavioral responses to these wavelengths^40^. Interestingly, mosquitoes show attraction during flight to patches of visual stimuli with long wavelengths present in human skin tones^41^. Thus, using red light could be a limitation that confounds the interpretation of optogenetic activation in mosquito host attraction assays. However, in our assays, genetic control mosquitoes showed minimal responses to broadly applied red light, even at strong intensities^8^ (**Figures 2-4**). We recommend genetic controls lacking either the driver or effector constructs and no-light controls to ensure behavioral effects are interpreted accurately. A control lacking the rhodopsin cofactor ATR is often used for optogenetics experiments in other species; however, the fish food used to feed mosquito larvae contains vitamin A, the precursor to ATR, so this is a less effective control in this system.

The optogenetic stimuli presented here are limited to continuous stimuli of adjustable light intensity. Optogenetic stimuli that closely recapitulate natural neuronal activity patterns are most likely to elicit the behavior of interest. Therefore, if neural recordings are available, the stimulus can be designed to match the observed intensity and temporal dynamics. When this is not known, the function of neurons can be screened using different stimuli and behavioral contexts, with the caveat that the results may not reflect the function of neurons in naturalistic behavior. Nevertheless, driving activity of neurons out of the natural range can be used to probe circuit or behavioral properties in unique ways just as electrophysiology can be used to probe cellular properties.

Currently, optogenetic tools have only been used in sensory neurons in the mosquito, an application that provides some advantages over the delivery of natural stimuli. Traditional strategies of measuring mosquito host seeking rely on the delivery of real CO_2_ gas. Using real CO_2_ requires air flow that has been filtered and humidified, which requires complex experimental setups to be able to precisely deliver and remove in short time increments^9,10,42,43^. Air flow and humidity are both cues for mosquito attraction and navigation, making optogenetic delivery appealing as a method to study the effects of olfactory stimuli separately from these other stimuli. Optogenetic tools have also previously been used to study the effect of sex and physiological state on behavior^8^, providing insight to these important questions in mosquito biology.

Apart from optogenetics, our assays make use of standard assays that have been used to assess mosquito behavior and come with some limitations. For ease of cleanup, the blood blanket makes use of an artificial meal that contains the only the tastants required for mosquitoes to engorge. This limitation could be overcome with the use of blood (as in the opto-membrane feeder) or biomimetic materials that recapitulate aspects of host tissue. Heat in the opto-thermocycler and blood blanket is delivered to walking or standing mosquitoes rather than flying mosquitoes and is limited by the ramp speed of the PCR thermocycler. Therefore, these assays are able to detect probing and engorgement behavior but not attraction. Another limitation of lab-based assays is that to understand the behavioral effect of specific cues, other cues that mosquitoes use are excluded. Increasingly complex lab assays and semi-field assays could be created to study additional aspects of the naturalistic environments in which mosquitoes feed on their hosts.

Another limitation of optogenetics in the mosquito is that few driver lines have been created beyond those for sensory neurons. To date, CO_2_ sensory neurons are the only neurons that driver lines have been developed for. Thus, its full use will be realized when it is combined with drivers that express in interneurons that cannot be directly stimulated by any other approach. Newly emerged resources such as the *Aedes aegypti* Mosquito Cell Atlas open the door for new advances in the field of mosquito neurogenetics^44^. Future work will be needed to develop such drivers for probing mosquito neural circuits. Similarly, additional optogenetic tools for inhibition will be needed to determine whether a neuron type is necessary for a particular behavior. Although here we demonstrate assays for host attraction and blood feeding, these approaches could be extended to study additional steps of host seeking and behaviors such as mating, oviposition, and nectar feeding. Optogenetic studies will provide new insight into how mosquito behavior is controlled at the neural circuit level and could lead to new approaches to prevent the spread of mosquito-borne diseases.

## Materials

| Name of Material | Equipment Company | Catalog Number | Comments/Description |
| --- | --- | --- | --- |
| AC Adapter | Jameco ReliaPro | 2197581 | Opto-membrane feeder structure |
| Acrylic Glue | United States Plastic Corp | 97562 | Opto-membrane feeder canister, blood blanket cage |
| Adenosine 5'-triphosphate disodium salt hydrate | Sigma Aldrich | A6419-1G | Artificial blood meal |
| All Trans-Retinal | Spectrum Chemical | R3041-1GM | Optogenetic rearing |
| Aluminum Breadboard | Thorlabs | MB12 | Opto-membrane feeder structure |
| Aluminum Sheet 0.0625 in | McMaster-Carr | 89015K37 | Blood blanket feeding plate |
| Aluminum Sheet 0.03125 in | McMaster-Carr | 89015K11 | Blood blanket feeding plate |
| Animal Part Tracker (APT) | Branson lab at HHMI Janelia Farm | N/A | Version 3.0, Opto-thermocycler reproduced data (Figure 2), Downloaded July 9, 2020 |
| Arduino IDE | Arduino | N/A | Version 1.8.12, used for stimulus control for opto-thermocycler, blood blanket, & opto-membrane feeder |
| Arduino Uno Rev3 Microcontroller Board | Arduino | A000066 | Opto-membrane feeder programming |
| Blackout Drape | Set Shop | 6520SF | Opto-thermocycler and blood blanket |
| Black Fiberglass Window Screen | Breakthrough Premium Products | IHLRS3684BL | Blood blanket cage |
| Black Tape | Thorlabs | T137-2.0 | Opto-thermocycler |
| Black Zip Ties | Cable Ties And More | ct242 | Opto-membrane feeder structure |
| Blue LED Lights | 21LEDUSA | RB5730BLWP | Mosquito rearing |
| Bottle | SKS Bottle & Packaging | 0604-07 | Opto-membrane feeder blood heater |
| Bottom Feeder Pellets | API | 5352983 | Feeding larvae |
| Camera | Blackfly | U3-13S2M-CS, FLI | Recording video for assays |
| Click Counter | ULine | H-7350 | Thinning larvae |
| CoolTerm | Roger Meier's Freeware | N/A | Version 2.0.1, used for serial port recording for opto-thermocycler, blood blanket, & opto-membrane feeder |
| Concentrated Bleach (8.25% Sodium Hypochlorite) | VWR | 89501-620 | Opto-membrane feeder cleaning |
| Computer | Asus | X1407Q | Running Assays |
| Defibrinated Sheep Blood | Hemostat Laboratories | DSB100 | Blood meal for opto-membrane feeder |
| Dome-topped hex 4-40 bolt | McMaster-Carr | 92949A105 | Blood blanket feeding plate |
| Epoxy | Loctite | 1363118 | Opto-thermocycler acrylic plate, Opto-membrane feeder structure |
| Ethanol | Decon Labs | 2716 | Sanitization |
| External Hard Drive | LaCie | LAC9000298 | Video Storage |
| ffmpeg | FFmpeg | N/A | Version 8.0.1, video pre-processing |
| Four Inch Cotton Wick | Richmond Dental | 201226 | Feeding adult mosquitoes |
| GitHub | GitHub, Inc | N/A | Repositories for code, part files, and SLEAP models for opto-thermocycler (v1.0.0, https://github.com/sorrellslab/opto-thermocycler/releases/tag/v1.0.0), blood blanket (v1.0.0, https://github.com/sorrellslab/bloodblanket/releases/tag/v1.0.0), and opto-membrane feeder (v1.0.0, https://github.com/sorrellslab/opto-membranefeeder/releases/tag/v1.0.0) |
| Gr3>CsChrimson | N/A | N/A | Created for this study, described in step 1.2 |
| Gr3-QF2 Mosquitoes | N/A | N/A | Genetic reagent, DOI: 10.1016/j.cell.2013.12.044 |
| ¼ Inch Black Acrylic | McMaster-Carr | 8560K259 | Opto-membrane feeder structure |
| ⅛ Inch Black Acrylic | McMaster-Carr | 8650K32 | Opto-membrane feeder structure |
| ⅛ Inch Clear Acrylic | McMaster-Carr | 8536K134 | Opto-membrane feeder canister |
| Incubator with Opaque Door | Caron | 7340-25 | Mosquito rearing |
| Infrared LED Light | Adafruit | 387 | Opto-thermocycler and blood blanket |
| Infrared LED Strip | Waveform Lighting | 7031.85 | Opto-thermocycler, blood blanket, and opto-membrane structure |
| Infrared Pass Filter | Edmund Optics | 65-796 | Recording video for assays |
| Insect Cage | Bugdorm | DP1000 | Adult mosquito enclosure |
| Janelia Automatic Animal Behavior Annotator (JAABA) | Branson lab at HHMI Janelia Farm | N/A | Opto-thermocycler reproduced data (Figure 2); Version 0.5.0 |
| Large Soup Cup | WebstaurantStore | 50016SOUPPLA | Mosquito rearing |
| LED Light | Luxeon Star | SP-01-D9 | Red light stimulus for Opto-thermocycler and blood blanket |
| LED Lens | Luxeon Star | 10209 | Opto-thermocycler and blood blanket |
| LED Lens Holder | Luxeon Star | 10235 | Opto-thermocycler and blood blanket |
| LED Lens Tape | Luxeon Star | LT-01 | Opto-thermocycler and blood blanket |
| LED Thermal Tape | Luxeon Star | LXT-S-12 | Opto-thermocycler and blood blanket |
| LED Heat Sink | Luxeon Star | N25-15B | Opto-thermocycler and blood blanket |
| Optical Construction Rails | Thorlabs | XE25L12, XE25L24, XE25L09, XE25T4, RA90, TRA6, TR12 | Opto-thermocycler and blood blanket structure |
| Optical Post | Thorlabs | TR12 | Opto-membrane feeder structure |
| Paper Towels | Pacific Blue | 27385 | Canister cleaning |
| Parafilm | Amcor | PM-996 | Blood blanket membrane, opto-membrane meal container |
| PCR thermocycler | Eppendorf | 2231001196 | Opto-thermocycler and blood blanket |
| Plastic Portion Cup | Choice | 999P2C | Feeding adult mosquitoes |
| Plastic Portion Cup Lid | Choice | 999PL2 | Feeding adult mosquitoes |
| Polycarbonate Pan | Cambro | 21410CWCHCL | Rearing larvae |
| Polycarbonate Pan Lid | Cambro | 10CWCH135 | Rearing larvae |
| Polycarbonate Pan Half Sized | Cambro | 24CW135 | Opto-membrane meal cleaning |
| Polycarbonate Pan Half Sized Lid | Cambro | 20CWCH135 | Opto-membrane meal cleaning |
| Polycarbonate Tube | McMaster-Carr | 8585K56 | Opto-membrane feeder canister |
| Power Meter | Coherent | 1299161 | All assays |
| QUAS-CsChrimson Mosquitoes | N/A | N/A | Genetic reagent, DOI: 10.7554/eLife.76663 |
| RGB Lights | Digikey | 289-1189-ND | Opto-membrane feeder stimulus delivery |
| Right Angle Clamp | Thorlabs | RA90 | Opto-membrane feeder structure |
| Silicone Gasket, 0.5 mm | McMaster-Carr | 1460N22 | Blood blanket feeding plate |
| SLEAP | Talmo Lab | N/A | Version 1.4.1a2, pose tracking of mosquitoes in each of the assays |
| Sodium Bicarbonate, 1M buffer soln., pH 8.0 | Thermo scientific | J62495.AP | Artificial blood meal |
| Sodium Chloride solution | Sigma-Aldrich | S5150-1L 100 3532552 | Artificial blood meal |
| Spray Bottle | Uline | S-11686 | Sanitization |
| Sucrose | Sigma Aldrich | S0389-500G | Feeding adult mosquitoes |
| Scotch Magic Tape | 3M | 104 | Opto-thermocycler and blood blanket acrylic lid |
| Thermocouple (Type T) | Harold G Schaevitz Industries LLC | CPTC-120-X-N | Opto-thermocycler and blood blanket |
| Thermocouple amplifier | Adafruit | MAX31856 | Opto-thermocycler and blood blanket |
| Timer | EnduraLight | RAC-MT | Controlling light cycle for mosquito rearing |
| Transfer Pipette | Thermo Scientific | SAM-225 | Thinning larvae |
| Trash Bag | Uline | S-23040 | Insect transportation |
| UV Resistant Mesh | McMaster-Carr | 87655K13 | Opto-membrane feeder canister |
| Water | Sigma Aldrich | W4502-1L | Artificial blood meal for blood blanket |
| Waxed Thread | McMaster-Carr | 6356K91 | Opto-membrane canister lid |
| Wire Mesh | McMaster-Carr | 9322T65 | Opto-membrane feeder structure |

## Acknowledgements

We thank Kim Lezon-Geyda, Fernanda Medeiros Contini, and Yaoyu Jiao for comments on the manuscript. This research was funded by NIH grant DP2AI177891. We thank the Neurotechnology Core funded by the Yale Kavli Institute for Neuroscience for technical advice. T.S. is an HHMI Freeman Hrabowski Scholar. This paper was typeset with the bioRxiv word template by @Chrelli: www.github.com/chrelli/bioRxiv-word-template

## References

1. UNICEF UNDP World Bank WHO Special Programme for Research. Global Vector Control Response 2017-2030. World Health Organization; 2017.

2. Knight KL, Stone A. A Catalog of the Mosquitoes of the World (Diptera: Culicidae). Entomological Society of America; 1959. doi:10.4182/umjb2446.1959.2

3. Raji JI, DeGennaro M. Genetic analysis of mosquito detection of humans. Curr Opin Insect Sci. 2017;20:34–38. doi:10.1016/j.cois.2017.03.003

4. Coutinho-Abreu IV, Riffell JA, Akbari OS. Human attractive cues and mosquito host-seeking behavior. Trends Parasitol. 2022;38(3):246–264. doi:10.1016/j.pt.2021.09.012

5. McMeniman CJ, Corfas RA, Matthews BJ, Ritchie SA, Vosshall LB. Multimodal integration of carbon dioxide and other sensory cues drives mosquito attraction to humans. Cell. 2014;156(5):1060–1071. doi:10.1016/j.cell.2013.12.044

6. Gillies MT. The role of carbon dioxide in host-finding by mosquitoes (Diptera: Culicidae): a review. Bulletin of Entomological Research. 1980;70(4):525–532. doi:10.1017/s0007485300007811

7. Geier M, Bosch O, Boeckh J. Influence of odour plume structure on upwind flight of mosquitoes towards hosts. The Journal of experimental biology. 1999;202 (Pt 12):1639–1648.

8. Sorrells TR, Pandey A, Rosas-Villegas A, Vosshall LB. A persistent behavioral state enables sustained predation of humans by mosquitoes. Elife. 2022;11:e76663. doi:10.7554/elife.76663

9. Dekker T, Geier M, Cardé RT. Carbon dioxide instantly sensitizes female yellow fever mosquitoes to human skin odours. Journal of Experimental Biology. 2005;208(15):2963–2972. doi:10.1242/jeb.01736

10. Breugel F van, Riffell J, Fairhall A, Dickinson MH. Mosquitoes Use Vision to Associate Odor Plumes with Thermal Targets. Current biology : CB. 2015;25(16):2123–2129. doi:10.1016/j.cub.2015.06.046

11. Liu MZ, Vosshall LB. General Visual and Contingent Thermal Cues Interact to Elicit Attraction in Female Aedes aegypti Mosquitoes. Curr Biol. 2019;29(13):2250–2257.e4. doi:10.1016/j.cub.2019.06.001

12. Chandel A, DeBeaubien NA, Ganguly A, et al. Thermal infrared directs host-seeking behaviour in Aedes aegypti mosquitoes. Nature. 2024;633(8030):615–623. doi:10.1038/s41586-024-07848-5

13. Tang R, Busby R, Laursen WJ, Keane GT, Garrity PA. Functional dissection of mosquito humidity sensing reveals distinct Dry and Moist Cell contributions to blood feeding and oviposition. Proc Natl Acad Sci. 2024;121(35):e2407394121. doi:10.1073/pnas.2407394121

14. Burgess L. Probing Behaviour of Ædes ægypti (L.) in Response to Heat and Moisture. Nature. 1959;184(4703):1968–1969. doi:10.1038/1841968a0

15. Hol FJ, Lambrechts L, Prakash M. BiteOscope, an open platform to study mosquito biting behavior. eLife. 2020;9:e56829. doi:10.7554/elife.56829

16. Martin-Martin I, Williams AE, Calvo E. Determination of Mosquito Probing and Feeding Time to Evaluate Mosquito Blood Feeding. Cold Spring Harb Protoc. 2023;2023(6):pdb.top107659. doi:10.1101/pdb.top107659

17. Boyd MF, Stratman-Thomas WK. Studies On Benign Tertian Malaria. 7. Some Observations on Inoculation and Onset*. Am J Epidemiology. 1934;20(2):488–495. doi:10.1093/oxfordjournals.aje.a118087

18. Swanson MM, Poodry CA. The shibire(ts) mutant of Drosophila: a probe for the study of embryonic development. Dev Biol. 1981;84(2):465–470. doi:10.1016/0012-1606(81)90416-4

19. Sweeney ST, Broadie K, Keane J, Niemann H, O’Kane CJ. Targeted expression of tetanus toxin light chain in Drosophila specifically eliminates synaptic transmission and causes behavioral defects. Neuron. 1995;14(2):341–351.

20. Baines RA, Uhler JP, Thompson A, Sweeney ST, Bate M. Altered Electrical Properties in Drosophila Neurons Developing without Synaptic Transmission. J Neurosci. 2001;21(5):1523–1531. doi:10.1523/jneurosci.21-05-01523.2001

21. Hamada FN, Rosenzweig M, Kang K, et al. An internal thermal sensor controlling temperature preference in Drosophila. Nature. 2008;454(7201):217– 220. doi:10.1038/nature07001

22. Corfas RA, Vosshall LB. The cation channel TRPA1 tunes mosquito thermotaxis to host temperatures. eLife. 2015;4. doi:10.7554/elife.11750

23. Greppi C, Laursen WJ, Budelli G, et al. Mosquito heat seeking is driven by an ancestral cooling receptor. Science (New York, NY). 2020;367(6478):681–684. doi:10.1126/science.aay9847

24. Allen WE, DeNardo LA, Chen MZ, et al. Thirst-associated preoptic neurons encode an aversive motivational drive. Science (New York, NY). 2017;357(6356):1149–1155. doi:10.1126/science.aan6747

25. Inagaki HK, Jung Y, Hoopfer ED, et al. Optogenetic control of Drosophila using a red-shifted channelrhodopsin reveals experience-dependent influences on courtship. Nature methods. 2014;11(3):325–332. doi:10.1038/nmeth.2765

26. Sahel JA, Boulanger-Scemama E, Pagot C, et al. Partial recovery of visual function in a blind patient after optogenetic therapy. Nat Med. 2021;27(7):1223–1229. doi:10.1038/s41591-021-01351-4

27. Klapoetke NC, Murata Y, Kim SS, et al. Independent optical excitation of distinct neural populations. Nature methods. 2014;11(3):338–346. doi:10.1038/nmeth.2836

28. Deisseroth K. Optogenetics. Nat Methods. 2011;8(1):26–29. doi:10.1038/nmeth.f.324

29. Potter CJ, Tasic B, Russler EV, Liang L, Luo L. The Q system: a repressible binary system for transgene expression, lineage tracing, and mosaic analysis. Cell. 2010;141(3):536–548. doi:10.1016/j.cell.2010.02.025

30. Herre M, Goldman OV, Lu TC, et al. Non-canonical odor coding in the mosquito. Cell. 2022;185(17):3104–3123.e28. doi:10.1016/j.cell.2022.07.024

31. Wohl MP, McMeniman CJ. Batch Rearing Aedes aegypti. Cold Spring Harb Protoc. 2022;2023(3):pdb.prot108017. doi:10.1101/pdb.prot108017

32. Hygiene AC of MEAS of TM and. Arthropod Containment Guidelines, Version 3.2. Vector Borne Zoonotic Dis. 2019;19(3):152–173. doi:10.1089/vbz.2018.2431

33. Pleil JD, Wallace MAG, Davis MD, Matty CM. The physics of human breathing: flow, timing, volume, and pressure parameters for normal, on-demand, and ventilator respiration. J Breath Res. 2021;15(4):042002. doi:10.1088/1752-7163/ac2589

34. Pereira TD, Tabris N, Matsliah A, et al. SLEAP: A deep learning system for multi-animal pose tracking. Nat Methods. 2022;19(4):486–495. doi:10.1038/s41592-022-01426-1

35. Mathis A, Mamidanna P, Cury KM, et al. DeepLabCut: markerless pose estimation of user-defined body parts with deep learning. Nature neuroscience. 2018;21(9):1281–1289. doi:10.1038/s41593-018-0209-y

36. Goodwin NL, Choong JJ, Hwang S, et al. Simple Behavioral Analysis (SimBA) as a platform for explainable machine learning in behavioral neuroscience. Nat Neurosci. 2024;27(7):1411–1424. doi:10.1038/s41593-024-01649-9

37. Tillmann JF, Hsu AI, Schwarz MK, Yttri EA. A-SOiD, an active-learning platform for expert-guided, data-efficient discovery of behavior. Nat Methods. 2024;21(4):703–711. doi:10.1038/s41592-024-02200-1

38. Skovorodnikov P, Zhao J, Buck F, et al. FERAL: A Video-Understanding System for Direct Video-to-Behavior Mapping. bioRxiv. Published online 2025:2025.11.16.688666. doi:10.1101/2025.11.16.688666

39. Kabra M, Robie AA, Rivera-Alba M, Branson S, Branson K. JAABA: interactive machine learning for automatic annotation of animal behavior. Nature methods. 2013;10(1):64–67. doi:10.1038/nmeth.2281

40. Autrum H, Bennett MF, Diehn B, et al. Comparative Physiology and Evolution of Vision in Invertebrates, A: Invertebrate Photoreceptors. 1979. doi:10.1007/978-3-642-66999-6_9

41. Alberto DAS, Rusch C, Zhan Y, Straw AD, Montell C, Riffell JA. The olfactory gating of visual preferences to human skin and visible spectra in mosquitoes. Nat Commun. 2022;13(1):555. doi:10.1038/s41467-022-28195-x

42. Dekker T, Cardé RT. Moment-to-moment flight manoeuvres of the female yellow fever mosquito (Aedes aegypti L.) in response to plumes of carbon dioxide and human skin odour. The Journal of experimental biology. 2011;214(Pt 20):3480–3494. doi:10.1242/jeb.055186

43. Demir M, Kadakia N, Anderson HD, Clark DA, Emonet T. Walking Drosophila navigate complex plumes using stochastic decisions biased by the timing of odor encounters. Elife. 2020;9:e57524. doi:10.7554/elife.57524

44. Goldman OV, DeFoe AE, Qi Y, et al. A single-nucleus transcriptomic atlas of the adult Aedes aegypti mosquito. Cell. 2025;188(25):7267–7290.e26. doi:10.1016/j.cell.2025.10.008

